# FGFR Signaling Coordinates the Balance Between Glycolysis and Fatty Acid β-Oxidation in Lymphatic Endothelial Cells

**DOI:** 10.1101/2020.05.20.107326

**Authors:** Hongyuan Song, Jie Zhu, Longhou Fang, Pengchun Yu

**Affiliations:** Cardiovascular Biology Research Program, Oklahoma Medical Research Foundation, Oklahoma City, OK 73104, USA; Center for Cardiovascular Regeneration, Department of Cardiovascular Sciences, Houston Methodist Research Institute, Houston, TX 77030, USA; Department of Obstetrics and Gynecology, Houston Methodist Institute for Academic Medicine, Houston Methodist Research Institute, Houston, TX 77030, USA; Department of Cardiothoracic Surgeries, Weill Cornell Medical College, Cornell University, New York City, NY 10065, USA

## Abstract

The balance between glycolysis and oxidative phosphorylation is believed to be critical for maintaining cellular bioenergetics, yet the regulation of such balance in lymphatic endothelial cells (LECs) remains unclear. Here we found that chemical inhibition of fibroblast growth factor receptor (FGFR) activity or knockdown of FGFR1, which has been shown to suppress glycolysis and consequently ATP production, induces substantial upregulation of fatty acid β-oxidation (FAO), but not glucose oxidation or glutamine oxidation in LECs. Mechanistically, blockade of FGFR-AKT signaling promotes the expression of CPT1A, a rate-limiting enzyme of FAO, in a PPARα-dependent manner. Metabolic analysis further showed that CPT1A depletion impairs ATP generation in FGFR1-deficient rather than wild-type LECs. This result suggests that FAO, which makes a minor contribution to cellular energy under normal conditions, can compensate for energy deficiency caused by FGFR inhibition. Consequently, CPT1A silencing potentiates the effect of FGFR1 knockdown on impeding LEC proliferation and migration. Collectively, our study identified an FGFR-regulated metabolic balance that is important for LEC growth.

## Introduction

Lymphatic vessels maintain tissue fluid homeostasis by absorbing excessive interstitial fluid and returning it to the venous circulation [1,2]. They are also involved in dietary fat absorption, immune cell trafficking, and several other physiological processes [1,2]. Congenital or acquired lymphatic defects may result in interstitial fluid retention and tissue swelling, a pathological condition termed lymphedema [3–5]. Assembly of the lymphatic network during murine development is a stepwise process [4,6]. A transcription factor prospero-related homeobox 1 (Prox1) starts to express in a subset of endothelial cells (ECs) of the anterior cardinal vein at embryonic day (E)9.5, and in turn, promotes the differentiation of the venous ECs to lymphatic ECs (LECs) [7,8]. This lymphatic fate specification process is followed by LEC sprouting out of the cardinal vein to form lymph sacs, which are a type of primitive lymphatic vascular structures, and further migration and proliferation of LECs to assemble into the highly-branched, mature lymphatic vascular plexus [9].

Lymphatic vascular development is driven by growth factor signaling [10]. Vascular endothelial growth factor (VEGF)-C and VEGF receptor 3, the cognate receptor highly expressed in LECs, have been shown by numerous studies to play an indispensable role in this process [11]. In contrast, the role of fibroblast growth factor (FGF) signaling in lymphatic vessel formation is much less understood. The FGF family of growth factors is composed of 22 members, with FGF2 as a robust mitogen to stimulate lymphangiogenesis both in vitro and in vivo [12,13]. The effects of FGFs are mediated by 4 types of FGF receptors (FGFRs, FGFR1 - FGFR4) [14]. FGFR1 is the predominant FGFR in LECs, and knockdown of FGFR1 is sufficient to impair LEC proliferation, migration, and tube formation induced by FGF2 in vitro [15]. However, during lymphatic vessel formation in vivo, the effect caused by loss of FGFR1 can be compensated by FGFR3. As such, we recently found that while genetic deletion of *Fgfr1* in LECs has no impact, double knockout of both *Fgfr1* and *Fgfr3* leads to profound defects of lymphatic vascular development [16]. Collectively, these results demonstrate an important but complex role played by FGFR signaling in lymphangiogenesis.

Recent advances have identified endothelial cell metabolism as a novel regulator of lymphatic vessel formation [17–20]. We and others found that glycolysis, which refers to a series of biochemical processes that convert glucose to pyruvate, provides more than 70% of ATP in LECs [21,16]. Our studies further show that the effect of FGFR signaling on lymphangiogenesis is controlled by glycolysis [16]. Knockdown of FGFR1—the most abundant member of the FGFR family—in LECs selectively reduces the expression of hexokinase 2 (HK2), and consequently suppresses glycolytic metabolism and ATP production, which are required for active angiogenic behaviors of LECs [16]. Functionally, lymphatic-specific deletion of *Hk2* in mice impairs lymphatic vascular development and abolishes the ability of FGF2 to promote lymphangiogenesis [16]. In addition to glycolysis, fatty acid β-oxidation (FAO) also has been shown to play a critical role in lymphatic vessel formation [22]. Carnitine palmitoyltransferase 1 (CPT1) is a group of rate-limiting enzymes of FAO, with CPT1A as the most abundant isoform in LECs [22]. Although FAO inhibition does not impair ATP generation under normal conditions, FAO is upregulated in LECs and controls lymphatic vessel growth through a histone acetylation-dependent mechanism [22]. Recent studies further demonstrate that glutamine metabolism is crucial for EC proliferation and angiogenesis by maintaining TCA cycle anaplerosis, asparagine synthesis, and redox balance [23,24]. Despite the importance of FAO and glutamine metabolism for EC growth, whether these metabolic pathways are regulated by FGFR signaling remains unknown.

In our current study, we sought to determine the impact of FGFR signaling inhibition on FAO, glutamine oxidation, and glucose oxidation. During this process, we unexpectedly discovered a balance between glycolysis and FAO, which plays an important role in regulating angiogenic behaviors of LECs.

## Results

### FGFR inhibition upregulates FAO while reducing glycolysis in LECs

Cellular energy in LECs is produced by glycolysis, glucose oxidation, glutamine oxidation, and FAO [21,16]. To assess the potential impact of FGFR signaling on the other metabolic processes that generate energy, we treated proliferating human dermal LECs (HDLECs) with 200 nM ASP5878, a highly specific FGFR inhibitor [25,26], for 2 days. HDLECs were cultured in EC growth medium and kept at subconfluency throughout the experiments to be maintained in the proliferative state. At the end of ASP5878 treatment, we incorporated 5-^3^H-glucose, 6-^14^C-glucose, ^14^C(U)-glutamine, or 9, 10-^3^H-palmitic acid into the culture medium and calculated flux of glycolysis, glucose oxidation, glutamine oxidation, and FAO through the measurement of ^3^H_2_O or ^14^CO_2_ generation. Consistent with our previous publication, ASP5878 treatment drastically reduced glycolytic flux (Fig. 1A). FGFR inhibition also impaired glucose oxidation but did not affect glutamine oxidation (Fig. 1B, C). However, in contrast to glycolysis and glucose oxidation that were both impaired by FGFR inhibition, FAO in HDLECs was significantly enhanced after ASP5878 treatment (Fig. 1D). To further confirm these observations with the FGFR chemical inhibitor, we transfected HDLECs with non-targeting (control) or FGFR1-specific siRNA. Three days following siRNA transfection, we trypsinized control or FGFR1 siRNA-transfected HDLECs and replated them into new culture dishes to obtain subconfluent cells the next day for analysis. Western blotting showed that FGFR1 proteins were effectively depleted by siRNA treatment (Fig. 1E, F). We further used metabolic flux analysis based on 5-^3^H-glucose and 9, 10-^3^H-palmitic acid to examine the effect of FGFR1 knockdown on glycolysis and FAO. Our data showed that FGFR1 siRNA treatment significantly upregulated FAO while reducing glycolysis (Fig. 1G, H). Together, our findings suggest the existence of a balance between glycolysis and FAO in LECs, as well as uncover the importance of FGFR signaling in this process.

**Figure 1:**
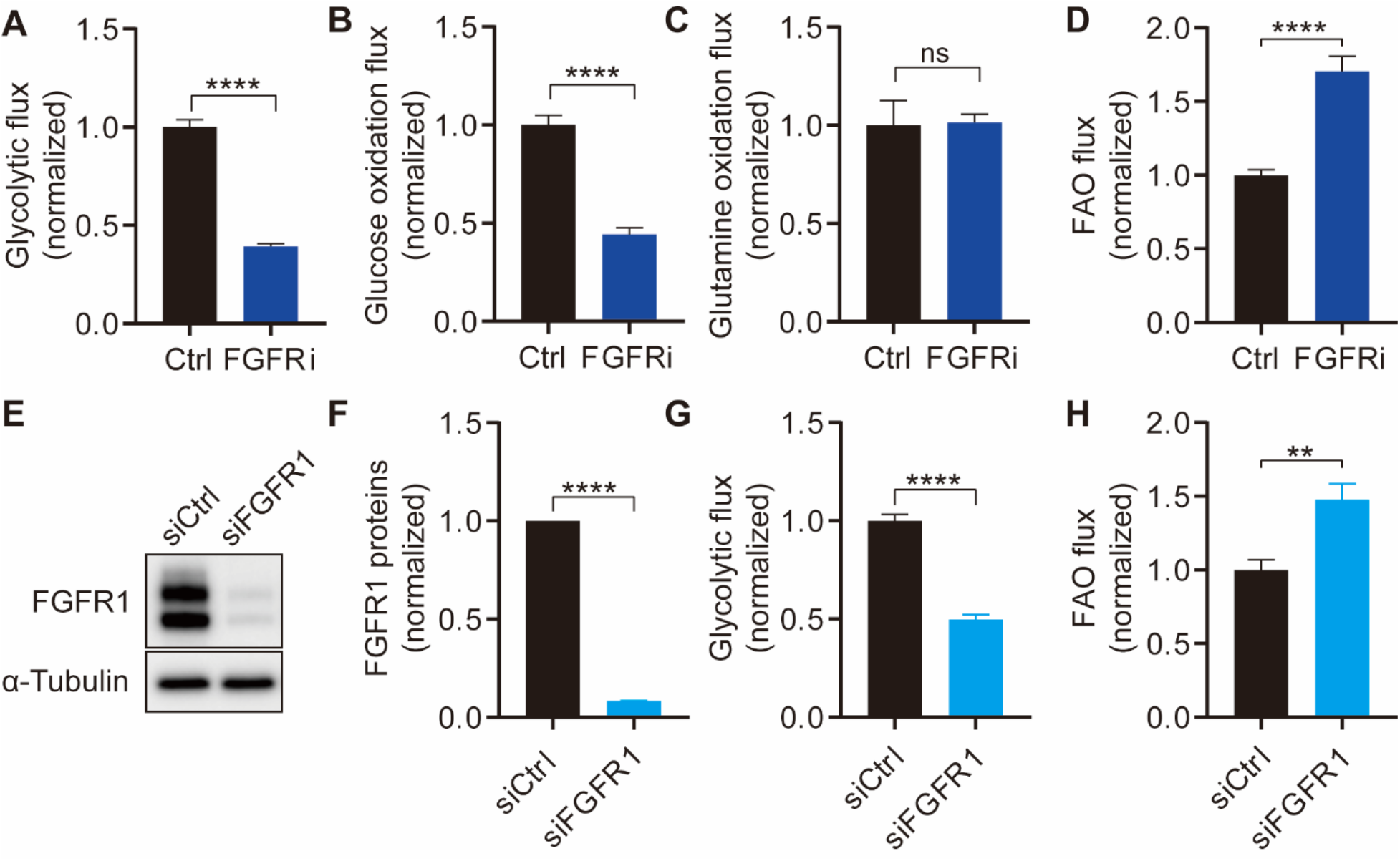
FGFR inhibition suppresses glycolysis but upregulates FAO in HDLECs. (**A-D**) Glycolytic flux (n=6) (**A**), glucose oxidation flux (n=4) (**B**), glutamine oxidation flux (n=4) (**C**), and FAO flux (n=6) (**D**) of HDLECs in the presence and absence of an FGFR inhibitor for 2 days. (**E, F**) Western blot analysis (**E**) and densitometric quantification (n=3) (**F**) of FGFR1 proteins in HDLECs transfected with non-targeting (Ctrl) or FGFR1-specific siRNA. (**G, H**) Glycolytic flux (n=6) (**G**) and FAO flux (n=6) (**H**) of HDLECs transfected with non-targeting (Ctrl) or FGFR1-specific siRNA. Data represent mean ± s.e.m., **p<0.01, ****p<0.0001, ns is not statistically significant, calculated by unpaired *t*-test.

### FGFR blockade increases CPT1A expression for promoting FAO in LECs

Next, we sought to elucidate the molecular mechanism by which FGFR signaling inhibition enhanced FAO. Because CPT1A promotes FAO as a rate-limiting step in LECs [22], we examined whether FGFR activity regulates CPT1A expression. Treatment of HDLECs with 200 nM ASP5878 for two days increased CPT1A protein levels while reducing the expression of HK2 (Fig. 2A-C), which is consistent with our previous report [16]. This data was confirmed by an experiment showing that FGFR1 knockdown led to reduction of HK2 but upregulation of CPT1A expression, albeit to a lesser extent than the FGFR inhibitor’s effect (Fig. 2D-G). We then assessed whether CPT1A elevation was required for the effect of FGFR inhibition on FAO. Our data showed that depletion of CPT1A expression by siRNA prevented ASP5878 or FGFR1 siRNA treatment from increasing FAO flux in HDLECs (Fig. 2H-J). Collectively, our findings suggest that FGFR signaling regulates FAO by controlling CPT1A expression.

**Figure 2:**
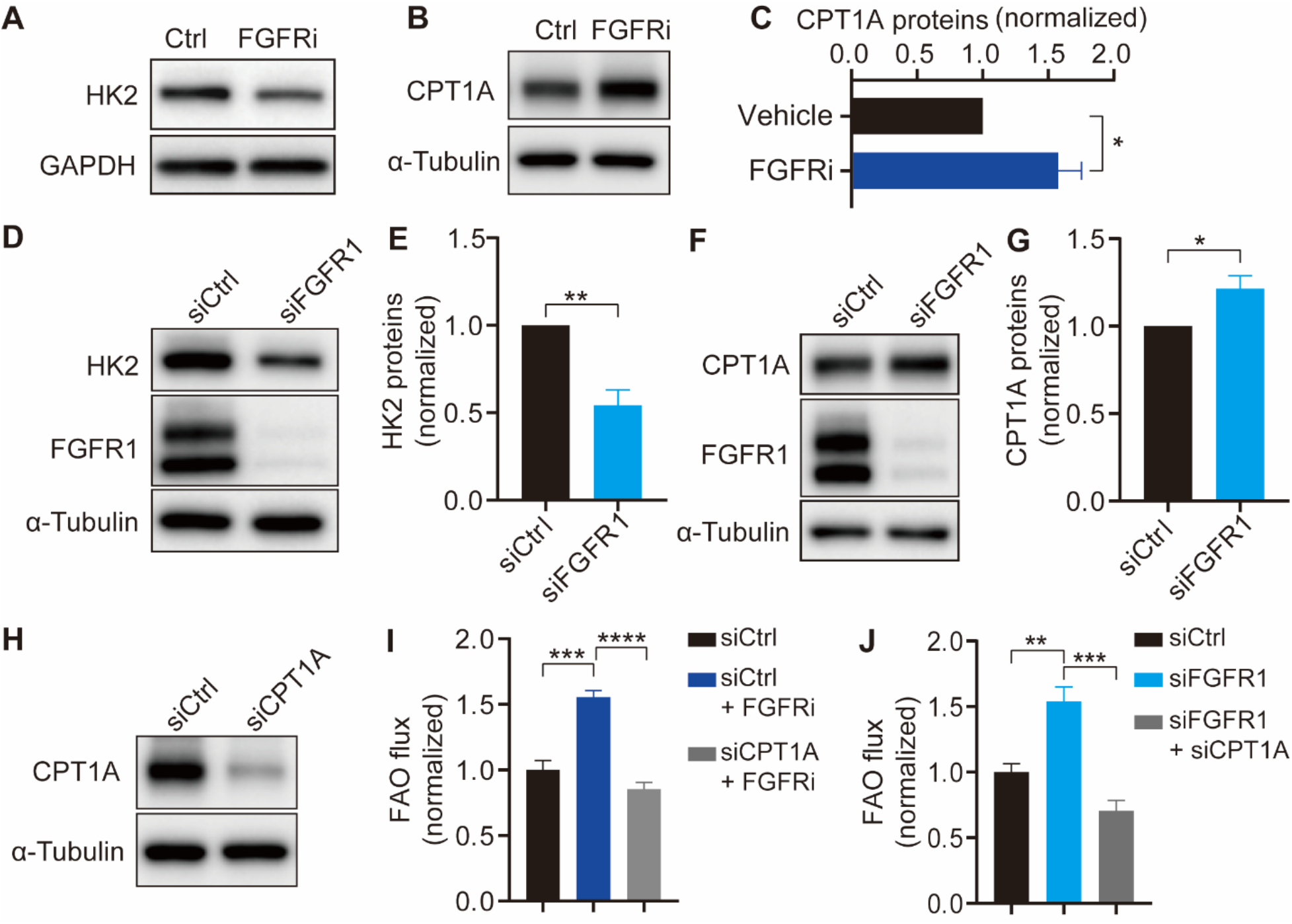
FGFR inhibition upregulates CPT1A expression for promoting FAO in HDLECs. (**A**) Western blot analysis of HK2 proteins in HDLECs in the presence and absence of an FGFR inhibitor for 2 days. (**B, C**) Western blot analysis (**B**) and densitometric quantification (n=3) (**C**) of CPT1A proteins in HDLECs in the presence and absence of an FGFR inhibitor for 2 days. (**D, E**) Western blot analysis (**D**) and densitometric quantification (n=3) (**E**) of HK2 proteins in HDLECs treated with non-targeting (Ctrl) or FGFR1 -specific siRNA for 4 days. (**F, G**) Western blot analysis (**F**) and densitometric quantification (n=3) (**G**) of CPT1A proteins in HDLECs treated with nontargeting (Ctrl) or FGFR1-specific siRNA for 4 days. (**H**) Western blot analysis CPT1A proteins in HDLECs treated with non-targeting (Ctrl) or CPT1A-specific siRNA for 4 days. (**I, J**) FAO flux of HDLECs with indicated treatments (n=4). Data represent mean ± s.e.m., *p<0.05, **p<0.01, ***p<0.001, ****p<0.0001, calculated by unpaired *t*-test (**C, E, G**) and ANOVA with Tukey’s multiple comparison test (**I, J**).

### FAO inhibition enhances the effect of FGFR signaling blockade on suppressing energy generation, proliferation, and migration of LECs

We next explored the functional significance of FAO upregulation for FGFR-deficient LECs. Previous studies show that FAO inhibition does not cause energy stress in ECs [27,22]. Consistently, we also found that CPT1A knockdown in HDLECs did not affect ATP production (Fig. 3A). However, CPT1A deficiency significantly potentiated the effect of FGFR1 siRNA on suppressing ATP production (Fig. 3A). Because proliferation and migration of LECs are highly energy-consuming, we then examined the impact of FAO inhibition on these cellular behaviors, which are critically involved in lymphatic vessel formation. Our data demonstrated that although knockdown of CPT1A or FGFR1 alone reduced LEC proliferation, their combined depletion resulted in more dramatic impairment (Fig. 3B). We further used an in vitro scratch assay to determine the role of FAO for LEC migration. We found that CPT1A knockdown did not affect migration when FGFR signaling was intact in LECs (Fig. 3C, D), which was consistent with a previous report [27]. However, CPT1 depletion aggravated migration defects caused by FGFR1 deficiency (Fig. 3C, D).

**Figure 3:**
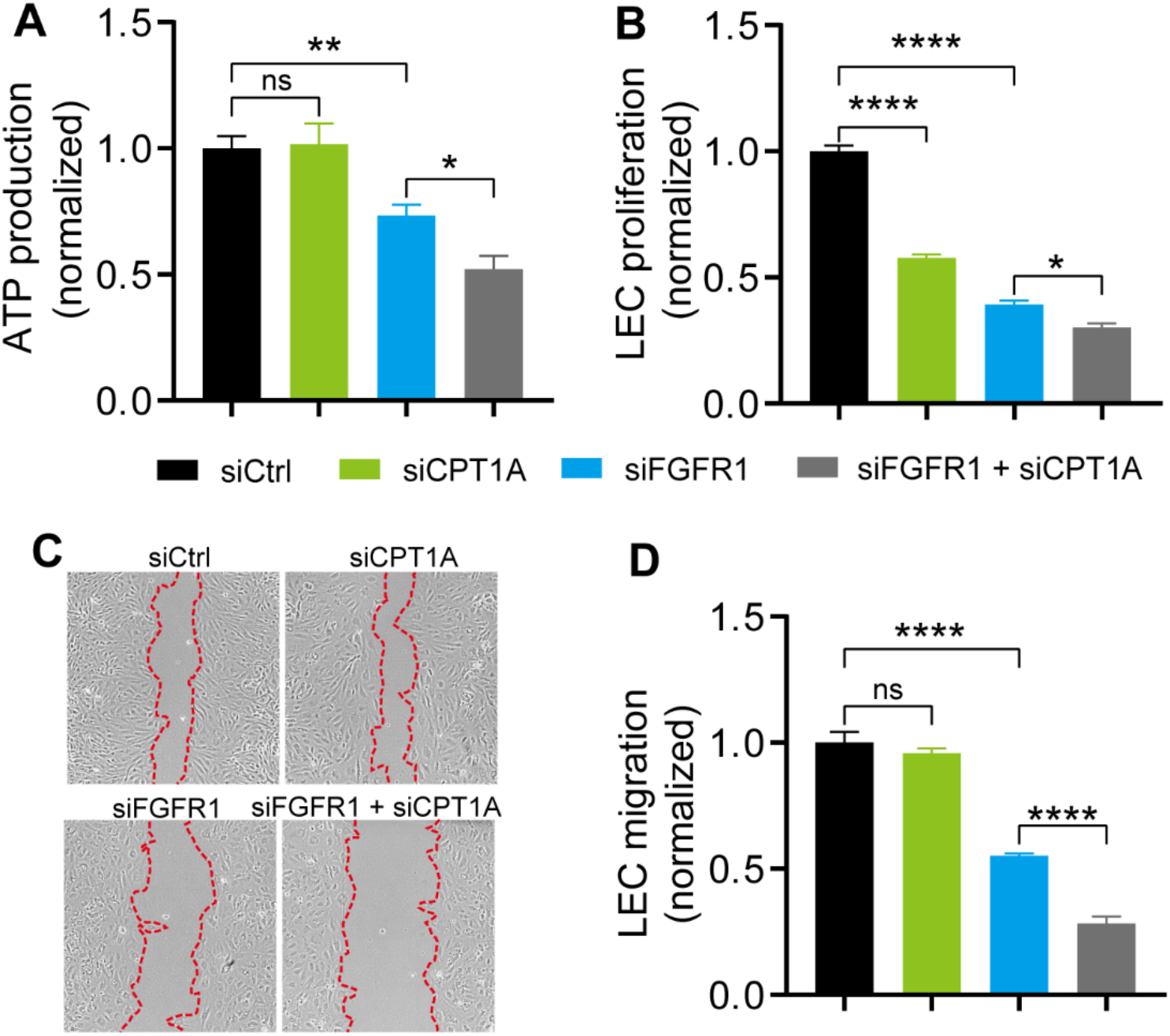
FAO inhibition potentiates the impact of FGFR blockade on suppressing ATP generation, proliferation, and migration of HDLECs. (**A**) ATP generation of HDLECs transfected with the indicated siRNA (n=4). (**B**) Proliferation of HDLECs transfected with the indicated siRNA (n=3). (**C**) Wound-healing assay to assess the migration of HDLECs transfected with siRNAs as indicated. Red dotted lines outline wound area at the last time points. (**D**) Quantification of migration area under different treatments (n=8). Data represent mean ± s.e.m., *p<0.05, **p<0.01, ****p<0.0001, ns is not statistically significant, calculated by ANOVA with Tukey’s multiple comparison test.

### Suppression of glycolysis per se does not lead to upregulation of CPT1A expression

To test whether CPT1A upregulation was due to FGFR inhibition-caused impairment of glycolysis, we treated HDLECs with 2-Deoxy-D-glucose (2DG), a potent inhibitor of glycolysis. Our data showed that 2DG failed to increase CPT1A levels (Supplemental Fig. 1). These data suggest that FGFR signaling controls CPT1A expression independent of glycolysis.

### Inhibition of AKT but not ERK signaling upregulates CPT1A expression and FAO

FGFR activity promotes phosphorylation and activation of downstream kinases, including AKT and ERK [28]. We sought to identify the downstream effectors that mediate the role of FGFRs in regulating CPT1A expression and FAO. We first knocked down AKT1 and AKT2, the major AKT isoforms expressed in ECs [29]. Our results showed that depletion of AKT 1 and AKT2 in HDLECs potently increased CPT1A levels and FAO flux (Fig. 4A-C). These effects were similar to what we observed in FGFR-deficient cells. In contrast, suppression of ERK signaling by downregulating ERK1 and ERK2, which are critical for arteriogenesis and integrity of the quiescent endothelium [30,31], had no impact on CPT1A expression and FAO (Fig. 4D-F). These data suggest that FGFR-dependent regulation of CPT1A and FAO is mediated by AKT rather than ERK signaling.

**Figure 4:**
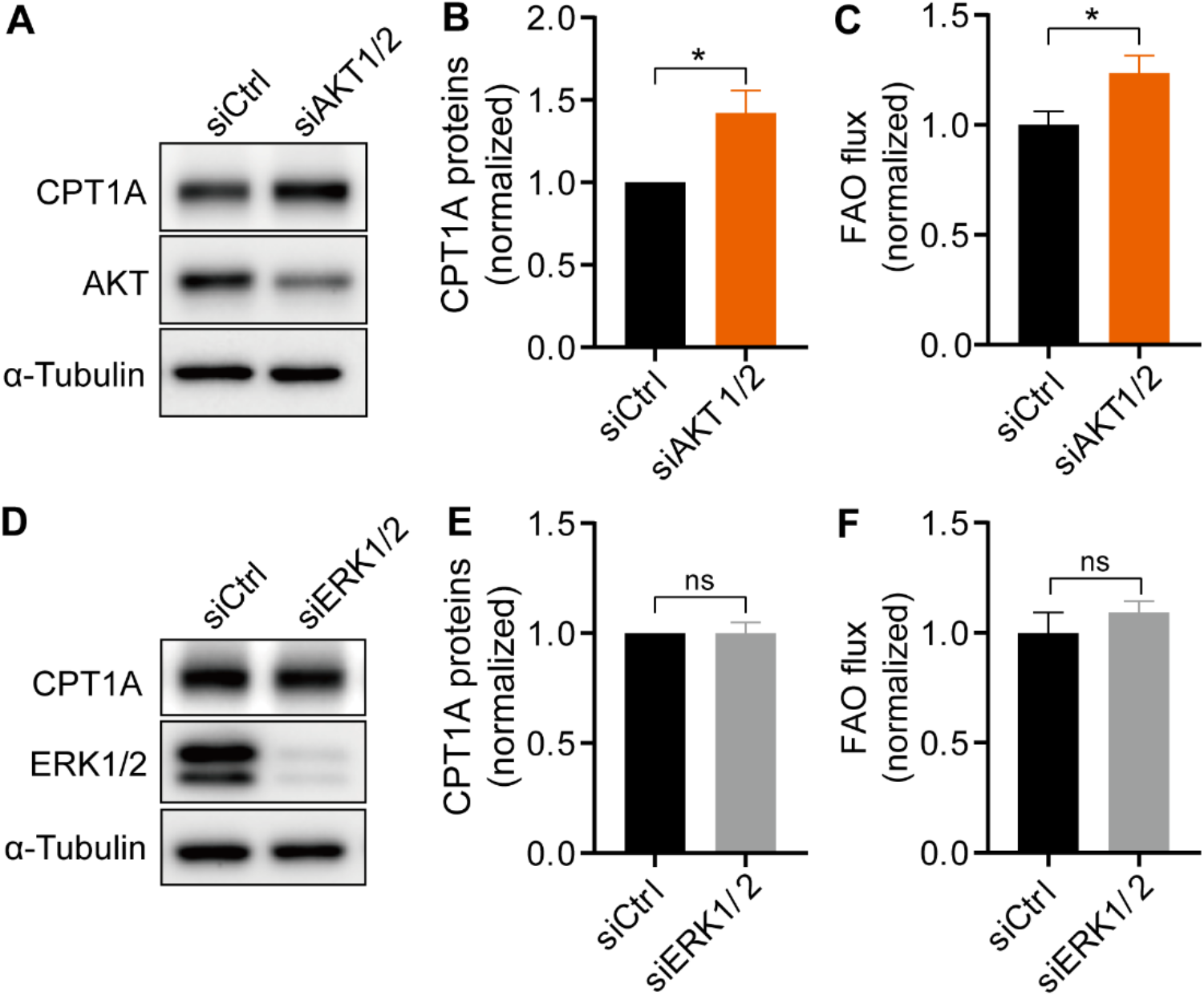
Inhibition of AKT but not ERK signaling upregulates CPT1A expression and FAO. (**A, B**) Western blot analysis (**A**) and densitometric quantification (n=3) (**B**) of CPT1A proteins in HDLECs treated with non-targeting (Ctrl) or AKT1/2-specific siRNA for 4 days. (**C**) FAO flux of HDLECs treated with non-targeting (Ctrl) or AKT1/2-specific siRNA for 4 days (n=6). (**D, E**) Western blot analysis (**D**) and densitometric quantification (n=3) (**E**) of CPT1A proteins in HDLECs treated with non-targeting (Ctrl) or ERK1/2-specific siRNA for 4 days. (**F**) FAO flux of HDLECs treated with non-targeting (Ctrl) or ERK1/2-specific siRNA for 4 days (n=4). Data represent mean ± s.e.m. (n=3), *p<0.05, ns is not statistically significant, calculated by unpaired *t*-test.

### Inhibition of FGFR-AKT signaling promotes CPT1A expression and FAO through PPARα

Our quantitative PCR (qPCR) experiments demonstrated that FGFR inhibition or FGFR1 knockdown significantly elevated *CPT1A* mRNA levels (Fig. 5A, B). Therefore, to further understand how FGFR signaling controls CPT1A expression, we sought to identify transcription factors that were involved in this process. Peroxisome proliferator-activated receptors (PPARs) are a family of nuclear receptor proteins that regulate transcription of genes involved in a variety of cellular processes, including those involved in lipid metabolism [32]. Recent studies have shown that PPARα and PPARγ, two PPAR isoforms, promote *CPT1A* transcription in hepatocytes and pulmonary arterial endothelial cells respectively [32,33]. These data led us to hypothesize that PPARα and PPARγ may mediate the effect of FGFR-AKT signaling on CPT1A expression. To test this idea, we used GW6471, a well-characterized PPARα inhibitor [34,35]. We found that treatment of HDLECs with GW6471 suppressed upregulation of CPT1A and FAO caused by the FGFR inhibitor ASP5878 (Fig. 5C-E). Similarly, inhibition of PPARα prevented FGFR1 knockdown-induced elevation of CPT1A expression and FAO flux (Fig. 5F-H). Our studies further showed that PPARα inhibition suppressed the effect of AKT1/2 siRNA on increasing CPT1A and FAO flux (Fig. 5I-K), suggesting that PPARα functions downstream of AKT in this process. In contrast to PPARα, inhibition of PPARγ by a specific inhibitor GW9662 [36,37], failed to block FGFR blockade-induced upregulation of CPT1A (Fig. 5L).

**Figure 5:**
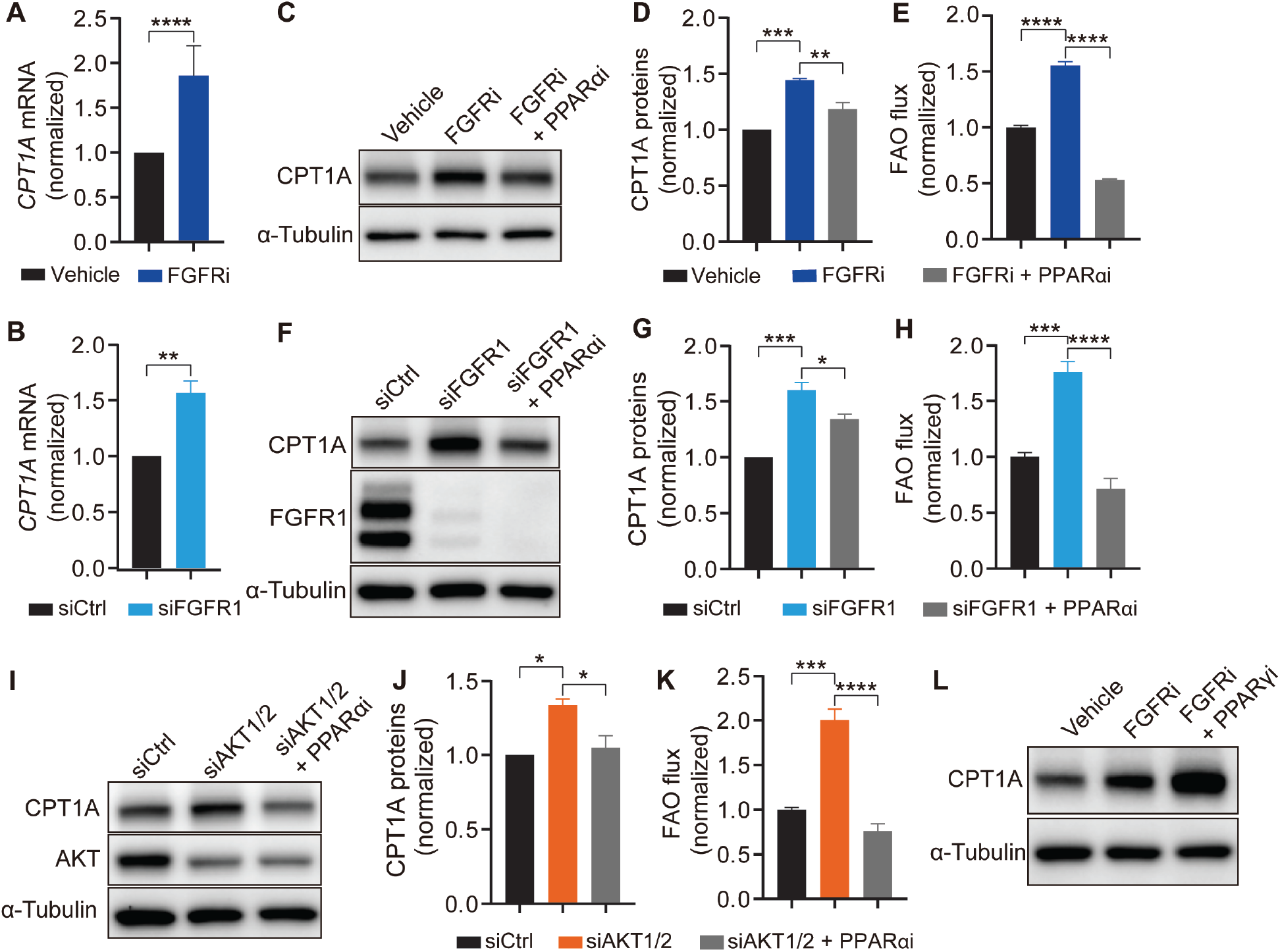
Inhibition of FGFR-AKT signaling upregulates CPT1A expression and FAO through PPARα. (**A**) qPCR analysis of *CPT1A* mRNA in HDLECs treated with vehicle or an FGFR inhibitor (n=3). (**B**) qPCR analysis of *CPT1A* mRNA in HDLECs transfected with nontargeting (Ctrl) or FGFR1-specific siRNA (n=3). (**C, D**) Western blot analysis (**C**) and densitometric quantification (n=3) (**D**) of CPT1A proteins in HDLECs treated with vehicle, an FGFR inhibitor, or a combination of the FGFR inhibitor and a PPARα inhibitor. (**E**) FAO flux of HDLECs treated with vehicle, an FGFR inhibitor, or a combination of the FGFR inhibitor and a PPARα inhibitor (n=4). (**F, G**) Western blot analysis (**F**) and densitometric quantification (n=3) (**G**) of CPT1A proteins in HDLECs treated with non-targeting (Ctrl) siRNA, FGFR1-specific siRNA, or both FGFR1-specific siRNA and a PPARα inhibitor. (**H**) FAO flux of HDLECs treated with non-targeting (Ctrl) siRNA, FGFR1-specific siRNA, or both FGFR1-specific siRNA and a PPARα inhibitor (n=4). (**I, J**) Western blot analysis (**I**) and densitometric quantification (n=3) (**J**) of CPT1A proteins in HDLECs treated with non-targeting (Ctrl) siRNA, AKT1/2-specific siRNA, or both AKT1/2-specific siRNA and a PPARα inhibitor. (**K**) FAO flux of HDLECs treated with non-targeting (Ctrl) siRNA, AKT1/2-specific siRNA, or both AKT1/2-specific siRNA and a PPARα inhibitor (n=4). (**L**) Western blot analysis of CPT1A proteins in HDLECs treated with vehicle, an FGFR inhibitor, or both the FGFR inhibitor and a PPARγ inhibitor. Data represent mean ± s.e.m. (n=3), *p<0.05, **p<0.01, ***p<0.001, ****p<0.0001, calculated by unpaired *t*-test.

## Discussion

In the current study, we uncovered an FGFR-dependent balance between glycolysis and FAO in LECs. While FGFR inhibition impairs glycolysis by reducing HK2 expression, it enhances FAO by upregulating CPT1A expression through AKT-PPARα signaling. Such metabolic balance allows FAO to compensate for energy deficiency caused by FGFR blockade. Functionally, suppression of FAO can exacerbate impaired LEC proliferation and migration caused by FGFR1 depletion. Our findings are reminiscent of the AMP-activated protein kinase (AMPK), which is an extensively studied energy sensor linking glycolysis with FAO. Energy stress due to defective glycolysis activates AMPK, which in turn enhances CPT1 activity and FAO by reducing the generation of a CPT1 inhibitor, malonyl-CoA [38]. However, we found that glycolytic inhibition caused by 2DG treatment, which activates AMPK in ECs [39], does not increase CPT1A levels (Supplemental Fig. 1), indicating that FGFR-mediated regulation of CPT1A expression is independent of AMPK. Therefore, our study may uncover a mechanism (promoting CPT1A expression) that functions in parallel with AMPK (enhancing CPT1 activity) to contribute to FGFR inhibition-induced FAO upregulation. Our results are also consistent with a recent study, which shows that compared to blood ECs (BECs) in the proliferative state, quiescent BECs, which were obtained through contact inhibition or activating Notch signaling, reduce glycolytic flux but potentiate FAO by increasing CPT1A [40]. In turn, the higher level of FAO sustains redox homeostasis but does not support energy production in quiescent BECs [40]. However, how CPT1A is induced in quiescent BECs remains to be determined.

Previous studies show that glycolysis generates most ATP in ECs, while FAO makes a minor contribution [21,16]. Consistently, CPT1A knockdown in BECs does not impair ATP production and migration, which is a highly energy-demanding process [27]. Although we obtained similar results in wild-type LECs, our data showed that CPT1A silencing decreases ATP levels and migration in FGFR1-deficient LECs (Fig 3A, C, D), suggesting that FGFR suppression makes LECs rely on FAO for acquiring energy. FAO has been demonstrated to regulate EC proliferation by contributing carbons for nucleotide synthesis [27,22]. As such, we found that knockdown of CPT1A is sufficient to impede LEC proliferation, but depletion of both CPT1A and FGFR1 caused a more dramatic effect (Fig. 3B), which likely reflects the outcome of dual inhibition of ATP generation and nucleotide synthesis.

AKT signaling is critical for cellular metabolism, especially in cancer cells [41]. A previous study shows that stimulation of murine prolymphocytic cells with interleukin 3 (IL-3), which is a hematopoietic growth factor, represses CPT1A expression and FAO [42]. Importantly, the effect of IL-3 is mediated by AKT [42]. Consistent with this report, we also found that depletion of both AKT1 and AKT2 upregulates CPT1A expression and FAO in HDLECs (Fig. 4), suggesting that AKT plays a conserved role in regulating FAO of different cell types. Although our data further showed that inhibition of FGFR signaling and AKT relies on PPARα to promote CPT1A expression, whether and how AKT regulates PPARα needs further investigation.

Our previous work demonstrates that chemical inhibition of FGFR activity reduces tumor lymphangiogenesis [16], which is critical for tumor metastasis [43]. Given the findings in the current study, targeting both FGFR signaling and FAO may offer a more effective means to suppress the formation of tumor-associated lymphatic vessels.

## Acknowledgments

We thank Ms. Summer Simeroth for proofreading the manuscript. This project was supported by AHA 19CDA34760260 and a grant from the Oklahoma Center for Adult Stem Cell Research to P. Y.

## Author Contributions

H.S. and J.Z. designed and performed the experiments and analyzed the data. L.F. provided critical suggestions and experimental materials for this project. P.Y. conceived the project, designed experiments, and supervised the studies. H.S. and P.Y. wrote the manuscript with the comments from the co-authors.

## Materials and Methods

### Cell culture and transfection

HDLECs (HMVEC-dLyNeo-Der Lym Endo EGM-2MV) were purchased from Lonza and cultured in EBM2 basal medium with EGM-2 MV BulletKit. HDLECs were tested negative for mycoplasma in Lonza. Culture medium was changed every other day. Tissue culture plates were coated with 0.1% gelatin (Sigma) for 30 minutes at 37 °C and washed with Dulbecco’s Phosphate-Buffered Saline (Life Technologies) before cell plating. FGFR1 siRNA, HK2 siRNA, CPT1A siRNA, ERK1 siRNA, ERK2 siRNA, AKT1 siRNA, AKT2 siRNA, and non-targeting siRNA were ordered from Qiagen and Dharmacon and transfected by Lipofectamine RNAimax reagent (Life Technologies).

To assay the effect of FGFR1, CPT1A, AKT, ERK, and HK2 knockdown on metabolic process and protein expression, HDLECs, transfected with relevant siRNA 3 days in advance, were replated and collected for indicated analysis approximately 16 hours later when the cell confluency reached ~80%. To assay the effect of PPARα inhibitor on metabolic process and protein expression after FGFR1 and AKT knockdown, HDLECs were transfected with indicated siRNAs 2 days in advance and then treated with PPARα inhibitor (10μM) for another 2 days. HDLECs were treated with FGFR inhibitor (200nM) for 2 days and then assayed the effect of FGFR inhibitor on metabolic process and protein expression. To assay the effect of PPARα inhibitor on the rescue of increased FAO and increased expression of CPT1A induced by FGFR inhibitor, HDLEC were treated with both FGFR inhibitor (200nM) and PPARα (10μM) inhibitor for 2 days and then assayed their effect on metabolic process and protein expression.

### Reagents

FGFR inhibitor (ASP5878) was purchased from Selleckchem (S6539). PPARα antagonist (GW6471) was purchased from Cayman chemical (11697). PPARγ antagonist was purchased from Sigma (M6191). 2-DG was ordered from Sigma (D8375). Luminescence ATP detection assay system (ATPlite) was purchased from PerkinElmer. For western blot analysis, the following antibodies were used: HK2 (Cell Signaling Technology, 2867), FGFR1 (Cell Signaling Technology, 9740), total-AKT (Cell Signaling Technology, 4691), ERK1/2 (Cell Signaling Technology, 4695), CPT1A (Abcam, 128568), GAPDH (Cell signaling Technology, 5174) and tubulin (Cell Signaling Technology, 2148). ImageJ was used for densitometry quantification of western blot bands.

### Measurement of glycolysis

Glycolytic flux was measured as previously described [44]. Briefly, subconfluent HDLECs cultured in 12-well plates were incubated with 1 ml per well EBM2 medium (containing appropriate amounts of serum and supplement) with [5-^3^H]-glucose (Perkin Elmer) for 2-3 hours. Then 0.8 ml per well medium was transferred into glass vials with hanging wells and filter papers soaked with H_2_O. After incubation in a cell culture incubator for at least 2 days to reach saturation, filter papers were taken out and the amount of evaporated ^3^H_2_O was measured in a scintillation counter.

### Measurement of FAO

FAO flux was measured essentially as reported [44]. Briefly, subconfluent HDLECs cultured in 12-well plates were incubated with 1 ml per well EBM2 medium (containing appropriate amounts of BSA, carnitine, and cold palmitic acid) with [9,10-^3^H]-palmitic acid (Perkin Elmer) for 6 hours. Then, 0.8 ml per well medium was transferred into glass vials with hanging wells and filter papers soaked with H_2_O. After incubation in a cell culture incubator for at least 2 days to reach saturation, filter papers were taken out and the amount of evaporated ^3^H_2_O was measured in a scintillation counter.

### Measurement of glucose oxidation

Glucose oxidation flux was measured as previously described [44]. Briefly, subconfluent HDLECs cultured in 12-well plates were incubated with 1 ml per well EBM2 medium (containing appropriate amounts of serum and supplement) with [6-^14^C]-glucose (Perkin Elmer) for 6 hours. Then, the cells were lysed using 12% perchloric (Sigma Aldrich, 244252) and the wells were covered immediately using filter papers soaked with hyamine hydroxide (Perkin Elmer, 2-19361). After incubation in a fume hood for at least 12 hours to reach saturation, filter papers were taken out and the amount of evaporated ^14^CO_2_ was measured in a scintillation counter.

### Measurement of glucose oxidation

Glutamine oxidation flux was measured as previously described [44]. Briefly, subconfluent HDLECs cultured in 6-well plates were incubated with 2 ml per well EBM2 medium (containing appropriate amounts of serum and supplement) with [^14^C(U)]-glutamine (Perkin Elmer) for 6 hours. Then, the cells were lysed using 12% perchloric (Sigma Aldrich, 244252) and the wells were covered immediately using filter papers soaked with hyamine hydroxide (Perkin Elmer, 2-19361). After incubation in a fume hood for at least 12 hours to reach saturation, filter papers were taken out and the amount of evaporated ^14^CO_2_ was measured in a scintillation counter.

### Western blotting

Cells were lysed using RIPA buffer and protein concentration was evaluated using bicinchoninic acid (BCA) assay. About 10 μg protein was loaded and separated by 10% SDS-PAGE, which was then transferred onto a polyvinylidene difluoride membrane (PVDF). The membrane was blocked with 5% non-fat milk for 45 minutes at room temperature and then incubated with relevant primary antibodies overnight at 4 degrees. After incubating with horseradish peroxidase-conjugated secondary antibodies, the membrane was visualized with super signal west pico chemoluminescent substrate using the ChemiDoc™ MP imaging system (Bio-Rad).

### Cell proliferation

About 80,000 HDLECs were seeded into each well of a 6-well plate and incubated overnight. Cells were transfected with control, FGFR1, CPT1A, or combined FGFR and CPT1A siRNA and cultured for 4 days. Then, cell numbers were counted using a hemocytometer (Hausser scientific Horsham).

### ATP production

ATP production was measured according to the manufacturer’s instruction (PerkinElmer, 6016943). Briefly, HDLECs, transfected with control, FGFR1, CPT1A, or combined FGFR and CPT1A siRNA 3 days in advance, were replated to a 96-well plate and ATP production analysis was performed approximately 12 hours later. 50 μL of mammalian cell lysis solution was added to each well and the plate was shaken for 5 minutes at 700 rpm. Then, 50 μL substrate solution was added to the cells and the plate was shaken for another 5 minutes at 700 rpm. After adapting the plate for 10 minutes, the luminescence was measured using FLUOstar Omega (BMG LABTECH).

### Wound-healing migration assay

HDLEC migration was measured in a wound-healing assay, which used ibidi Culture-Inserts (ibidi) to generate the wound. An ibidi Culture-Insert has dimensions 9 mm × 9 mm × 5 mm (width × length × height) and is composed of two wells. One or two inserts were placed into one well of six-well plates. HDLECs, transfected with control, FGFR1, CPT1A, or combined FGFR and CPT1A siRNA 3 days in advance, were replated to the wells of the inserts. When cells became fully confluent after attachment, Culture-Inserts were carefully removed by sterile tweezers to start cell migration. For studying the effect of FGFR1 and CPT1A siRNA on migration, the wound-healing process was monitored for approximately 11 hours. A Nikon ELIPSE Ti microscope with an ANDOR camera was used to image cells at the first time point (*T*_0_) and the last time point (*T*_end point_). For data analysis, ImageJ was used to measure the wound area at *T*_0_ and *T*_end point_. Migration area was obtained by subtracting area at *T*_end point_ from area at *T*_0_.

### qPCR analysis

Total RNA was extracted from HDLECs using the TRIzol reagent (ThermoFisher) according to the manufacturer’s instructions. The concentration and quality of RNA were analyzed by Nanophotometer (IMPLEN). cDNA synthesis was performed using the iScript cDNA synthesis kit (Bio-rad). qPCR was performed with Evagreen qPCR Master Mix (Bullseye) using the CFX96 Real-Time system (Bio-rad).

### Statistical analysis

Statistical significance between two groups was determined by two-tailed unpaired *t*-test (assuming normal distribution), and statistical significance between multiple groups was calculated using one-way ANOVA with post-hoc tests. Data represent mean ± s.e.m.

**Supplemental Figure 1:**
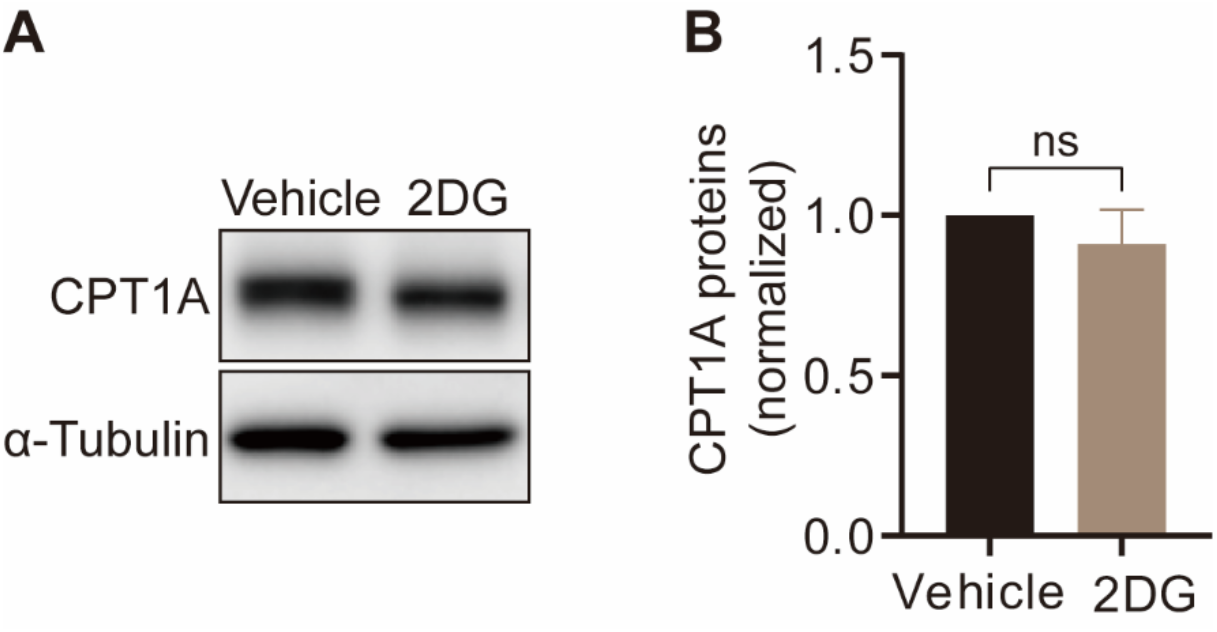
Inhibition of glycolysis per se does not affect CPT1A expression. (**A, B**) Western blot analysis (**A**) and densitometric quantification (**B**) of CPT1A proteins in HDLECs treated with vehicle or 2DG. Data represent mean ± s.e.m. (n=3), ns is not statistically significant, calculated by unpaired *t*-test.

